# LED Zappelin’: An open source LED controller for arbitrary spectrum visual stimulation and optogenetics during 2-photon imaging

**DOI:** 10.1101/2020.03.10.986000

**Authors:** M.J.Y. Zimmermann, A.M. Chagas, P. Bartel, S. Pop, L.L. Prieto Godino, T. Baden

## Abstract

Two-photon (2P) microscopy is a cornerstone technique in neuroscience research. However, combining 2P imaging with spectrally arbitrary light stimulation can be challenging due to crosstalk between stimulation light and fluorescence detection. To overcome this limitation, we present a simple and low-cost electronic solution based on an ESP32 microcontroller and a TLC5947 LED driver to rapidly time-interleave stimulation and detection epochs during scans. Implemented for less than $100, our design can independently drive up to 24 arbitrary spectrum LEDs to meet user requirements. We demonstrate the utility of our stimulator for colour vision experiments on the *in vivo* tetrachromatic zebrafish retina and for optogenetic circuit mapping in *Drosophila*.

https://github.com/BadenLab/LED-Zappelin

## 1. Introduction

Advances in fluorescence microscopy techniques and the development of genetically encoded biosensors (Akerboom et al., 2012; Marvin et al., 2013) have allowed interrogating the structures and functions of biological systems at ever increasing depth (Mostany et al., 2014; Svoboda and Yasuda, 2006). In particular, two-photon (2P) imaging has enabled high temporal and spatial resolution imaging deep within intact tissues due to a low degree light scattering of the infrared laser (Helmchen and Denk, 2005).

And yet, combining 2P imaging with additional light stimulation – for example for visual stimulation or for driving optogenetic actuators – has remained challenging because the stimulation light can interfere with fluorescence detection. This can result in light artefacts in the image and/or may damage sensitive fluorescence detection equipment (e.g. photomultiplier tubes, PMTs). A temporal separation between light stimulation and fluorescence detection, for example during the scan-retrace, can ameliorate these problems (Euler et al., 2019a).

To limit flickering artefacts, the rate of interweaving stimulation and imaging epochs should ideally be substantially beyond the integration time of the to-be stimulated system. Accordingly, line-synchronisation (typically 500 – 1,000 Hz) is usually preferred over frame-synchronisation (typically in the order of 10s of Hz). For this, both mechanical and electronic solutions are possible. For example, some systems rely on an optical chopper and/or mechanical shutter with microsecond performance for rapid gating of a constant light source (Alfonso-Garcia et al., 2019; Yang et al., 2019). However, those systems tend to be bulky and/or expensive and can introduce mechanical vibrations. Alternatively, the problem can be readily solved electronically, for example through use of a microcontroller. Here, we present such a solution. Our system can line-synch up to 24 independent LED channels, and can be assembled from off-the-shelf components for substantially below $100. This provides for flexible options of spectrally diverse light stimulation during 2-photon scanning and comfortably provides sufficient power to drive standard optogenetics actuators such as CsChrimson (Klapoetke et al., 2014). Alongside, we also provide a custom 3D-printed casing, design suggestions for optically combining LED banks using Thorlabs parts, and an alternative 3D-printed LED holder and microscope chamber. For software control, we provide custom Arduino scripts to flexibly programme stimulation protocols.

## 2. Hardware description

### 2.1. Overview

The challenge when combining 2P-microscopy and light stimulation centres around the potential spectral overlap of photodetector (e.g. photomultiplier) sensitivities and the stimulus light. Spectral separation is not always enough depending on the required excitation light, the available fluorescent probes or limitations in the microscope’s optical design and its components (e.g. spectral filter imperfections or reflections). In this case, temporal separation between stimulation and fluorescence recording can be helpful in reducing crosstalk (Euler et al., 2019a). This requires time-precise control over the stimulus light, and a means to synchronise this control with a readout of the scan pattern.

High performance LEDs, unlike conventional light sources, near instantly illuminate at full intensity when current is applied. They emit narrow-spectrum light and can be switched on and off within nanoseconds while providing high emission stability (Hohman, 2007). Similarly, even basic microcontrollers provide for suitable time resolution to control LED state in the microsecond range. We therefore built a microcontroller and LED-based stimulator system to interweave the laser excitation timing with LED illumination: By turning off the LEDs during the laser scanning period and turning them on during the mirror retrace period, a clean separation between stimulation and emission lights can be achieved while nevertheless delivering sufficient average light to stimulate photo-sensitive cells and commonly used optogenetic actuators.

For synchronisation with the imaging system, we use a “blanking” signal from the scan-software: Most conventional 2P-imaging software solutions can send a transistor-transistor-logic (TTL) signal which runs synchronised to the fast scanning mirrors through their digital acquisition (DAQ) system. Such a TTL signal is for example frequently used to synchronise a Pockels cell for rapid regulation of laser power during the retrace period of a raster scan to avoid phototoxicity (Icha et al., 2017). We use the same signal to synchronise the LED illumination with the retrace period (Fig. 1d) (Euler et al., 2019a; Franke et al., 2019). This strategy works for both galvo-galvo and the substantially faster line rates of resonant scanning systems.

**Figure 1.**
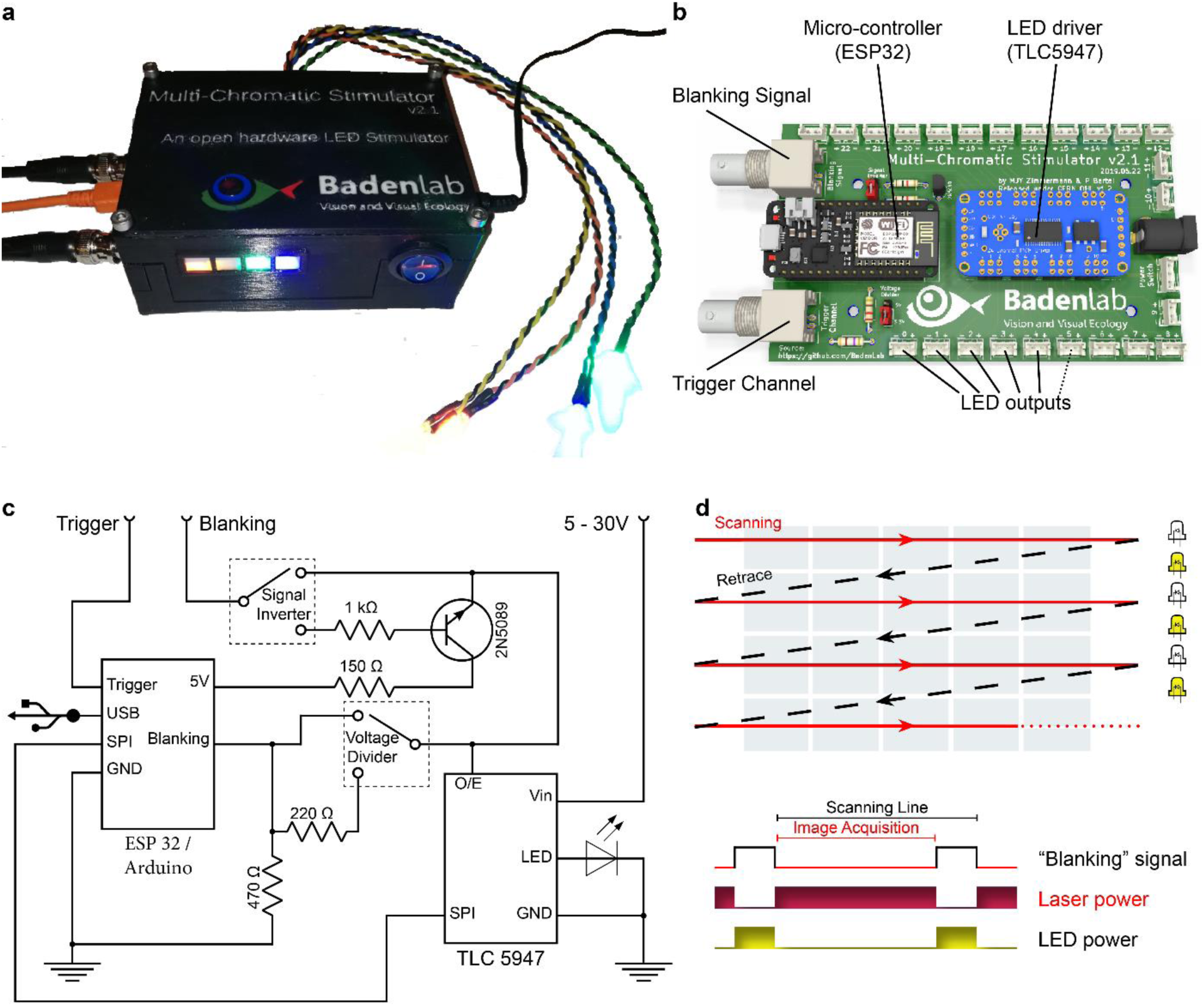
Stimulator design. **a.** A fully assembled stimulator. **b.** Rendering of the custom-printed circuit board which accommodates the microcontroller, the LED driver and up to 24 LED channels. **c.** Schematics illustrating the circuit that controls the LED output. The blanking input can be inverted by a switch before reaching the *output enable* pin on the LED driver (electronically switching off the LEDs) and sending the signal to the micro-controller. A second switch controls the blanking signal voltage as it needs to be adapted depending on the logic of the microcontroller used (3.3 V for ESP32, 5 V for Arduino). The microcontroller controls the LED driver through an SPI connection and sends a trigger signal output to an external DAQ-system. KiCAD schematic are available on the GitHub repository. **d.** Illustration of the raster scan method described. The “blanking signal” is synchronous with the scanning logic, enabling the LEDs during the scanning mirrors retrace (black) and shutting them off during the acquisition (red), therefore providing temporal separation between stimulation and detection (schematic in (d) inspired from (Euler et al., 2019a)).

### 2.2 Electronics

The stimulator consists of an ESP32 micro-controller (Adafruit ESP32 feather, built around the ESP32 system on a chip, Espressif), an LED driver (Adafruit TLC5947, built around the TLC5947 chip, Texas instruments), and common off-the shelf electronics components (Fig. 1b). Designed around a custom-built printed circuit board (PCB) (Fig. 1b), the stimulator assembly is intuitive and does not require previous electronics nor soldering experience (c.f. Supplementary video 1). The total cost for electronic parts is currently below $100.

The ESP32 is a low cost and relatively recent (released in Sept. 2016) type of microcontroller operating at 240MHz which provides enough processing performance to control the light output in the kHz range. The microcontroller, through Serial Peripheral Interface (SPI), communicates with the LED driver, and allows for independent control of up to 24 LED channels, each with 12 bits resolution (4,096 “grey levels”). As one example use-case, our visual stimulator for zebrafish retina experiments uses four spectrally distinct stimulus LEDs plus another set of 4 “proxy”-LEDs which can be embedded in the 3D-printed chassis as a visual control for the experimenter (Fig. 1a).

### 2.3 LED control and linear modulation

One of the key features of the chosen LED driver is its blanking input (labelled O/E on the TLC5947 board for “output enable”), which allows for the fast and simultaneous switching of all LEDs. When the blanking input is high, all 24 current outputs are forced off. When the blanking input is low, all constant current outputs are controlled by the grayscale Pulse-width-modulation (PWM) timing controller which is reset during the blanking, thus providing a stable light output (Fig. 2a).

**Figure 2.**
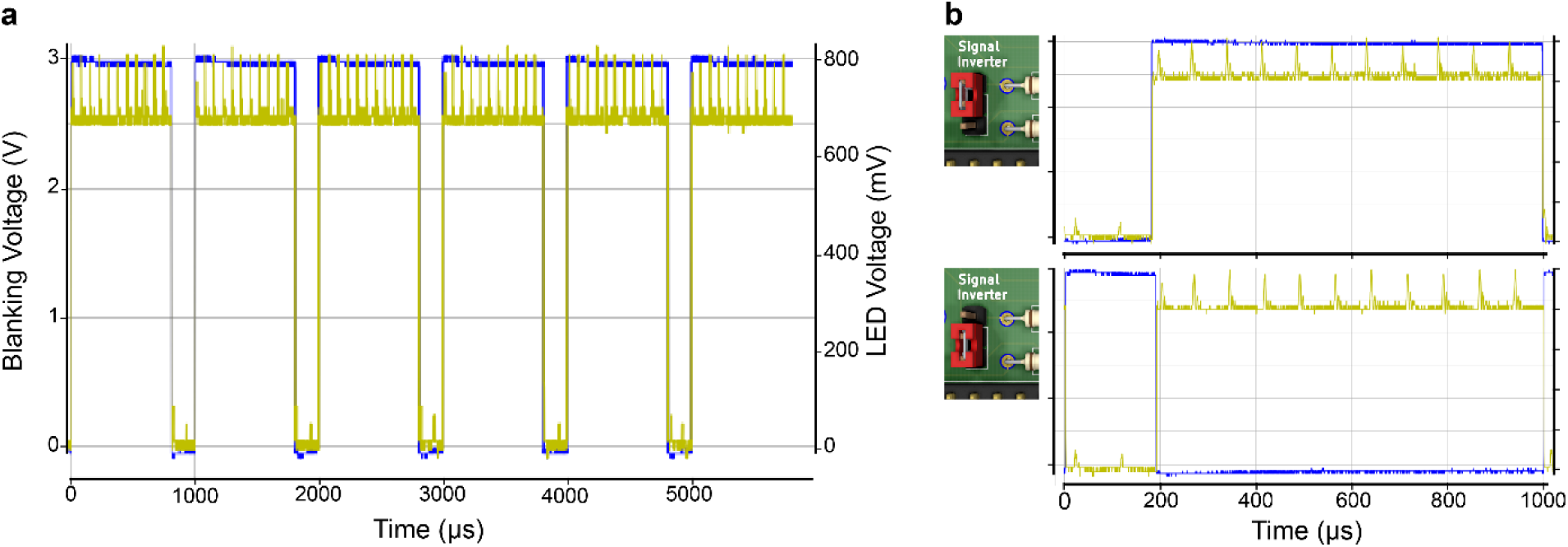
The blanking signal controls LED illumination. **a.** Oscilloscope reading of the blanking signal (blue) efficiently switching off an LED (yellow). The blanking is operated here without noticeable delay. **b.** as **a.**, shown for a 1 ms scanning cycle, the two possible configurations for the blanking signal input, with a *LOW* (top) and a *HIGH* (bottom) blanking signal input for an inverted and original signal input, respectively.

The blanking signal also serves as the stimulator’s external clock: the stimulator counts the number of TTL pulses (i.e. scan-lines) and from here computes the accurate timing to move through a pre-programmed stimulus sequence (c.f. 6.2).

For experiments in vision neuroscience, we use a custom movable-objective microscope (MOM)-type 2P system (Euler et al., 2009) controlled by “ScanM” (developed by W. Denk, M. Müller and T. Euler). ScanM is configured to provide a HIGH signal during blanking which can be fed directly to the TLC5947 (Euler et al., 2019b). However, in our example optogenetics experiment, we use another software package (ScanImage (Pologruto et al., 2003)) on a custom-made 2P microscope (Independent NeuroScience Services), which instead provides a LOW signal during the retrace. We therefore incorporated a signal inverter which can be enabled through a jumper (Fig. 2b). When the jumper is close to the ESP32, the signal goes straight to the TLC5947, while when it is placed away from the microcontroller, the signal is inverted by passing a logical NOT gate (Fig. 2b).

This option thus offers the possibility for a single design to be easily adapted to different software systems.

Another important feature to consider when using a constant-current sink LED driver like the TLC5947 is its ability to drive a near linear LED output. Indeed, an LED brightness modulated directly by a microcontroller does not necessarily scale linearly with pulse width but rather adopts a sigmoid dependency (Fig. 3). The use of an adequate LED driver therefore does not require further gamma correction from the experimenter.

**Figure 3.**
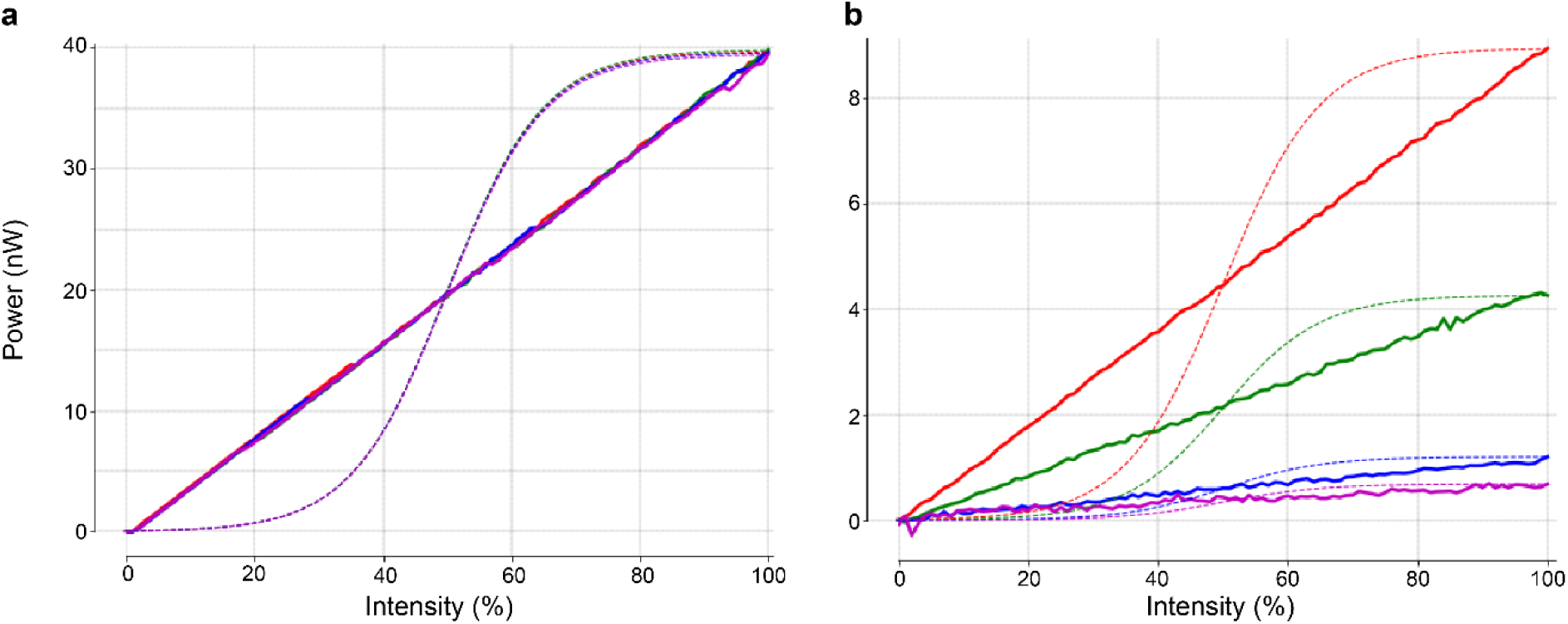
LED electrical power over duty cycle (PWM). **a.** Power recording of a 4 LED system using the TLC5947 (solid lines) and their expected brightness if directly controlled by a microcontroller (dashed lines). All LEDs have been set up to the same power (40 nW), with equal maximal intensity values in the Arduino code (c.f. 6.3). **b.** as **a.** but with LEDs set up at different maximal intensities in the Arduino code. Here the linearity of the LED intensity output remains constant.

Furthermore, the use of a dedicated constant current LED driver tends to improve LED stability over time as well as its life span. Such a driver ensures that the current drawn by the LED does not lead to thermal runaway which can cause irreversible damage. This is particularly important for short wavelength LEDs which tend to decay rapidly as they usually require higher power supply leading to higher thermal runaway. This characteristic thus necessitates regular recalibration or replacement.

### 2.4 A trigger signal for synchronising stimulation protocols and image acquisition

The stimulator is equipped with an output trigger channel that can be connected to the 2P setup’s DAQ-system to precisely time-align the light stimulation with imaging data. By default, this is a 5 V TTL trigger-signal of 25 ms duration which starts as soon as a stimulus sequence begins via software control (see below) and is thereafter repeated with a pulse of exactly 1 Hz (1 trigger every 1,000 ms). These numbers are easily adjusted in the Arduino code provided. Due to the ESP32’s high processing speed, this trigger signal is reliable and precise within 0.1 μs (Fig. 8e) (see below).

**Figure 4.**
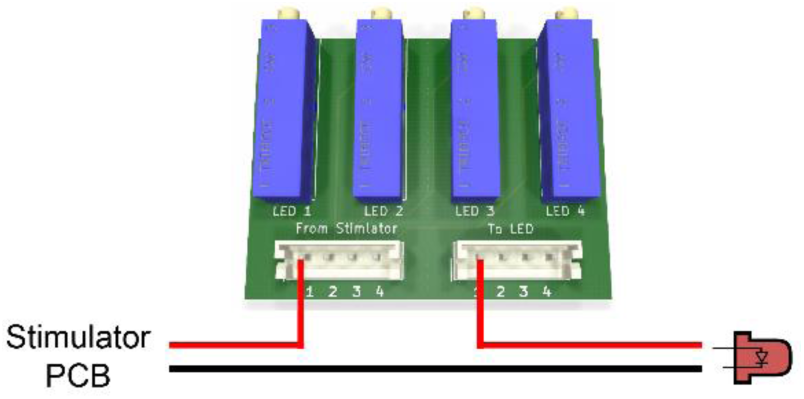
Potentiometer Mount PCB. Wiring example of the LED channel 1 to its trimmer potentiometer. Note that LED polarity as indicated on the stimulator PCB must be respected.

**Figure 5.**
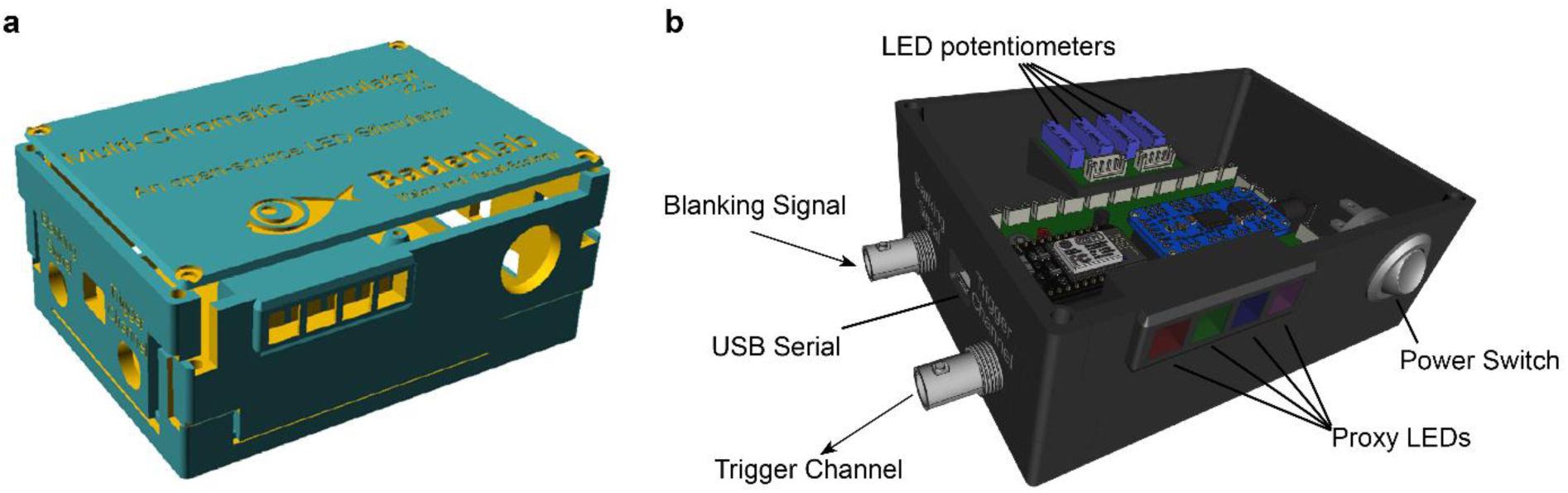
Stimulator box. **a.** Rendering of the stimulator box 3D files set here by default for 4 LED channels and 4 proxy LEDs. **b.** Rendering of the fully mounted stimulator with all PCBs and components tightly fitting their respective space.

**Figure 6.**
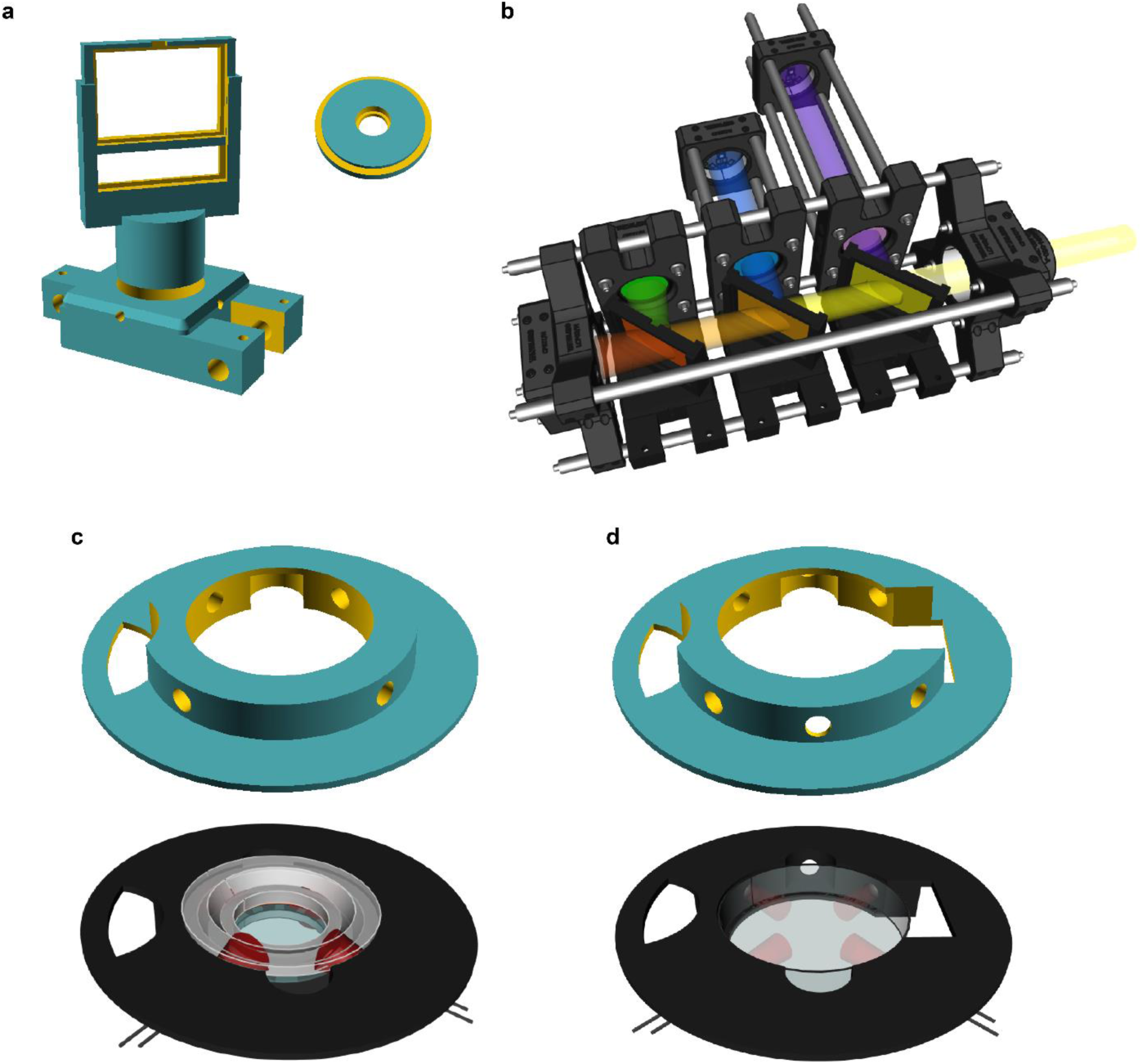
3D-printed Illumination systems. **a.** SCAD files for adapting 5mm LEDs and dichroic mirrors to standard 30mm optomechanical system. **b.** Rendering of the LED illumination system for the visual experiment. **c.** For optogenetics experiments, we designed a mounting platform that holds four 5mm LEDs and can fit a RC-40HP chamber (SmartEphys, Warner Instrument). **d.** Same as **c.** but designed to fit a small petri dish (ø 35mm) lid.

**Figure 7.**
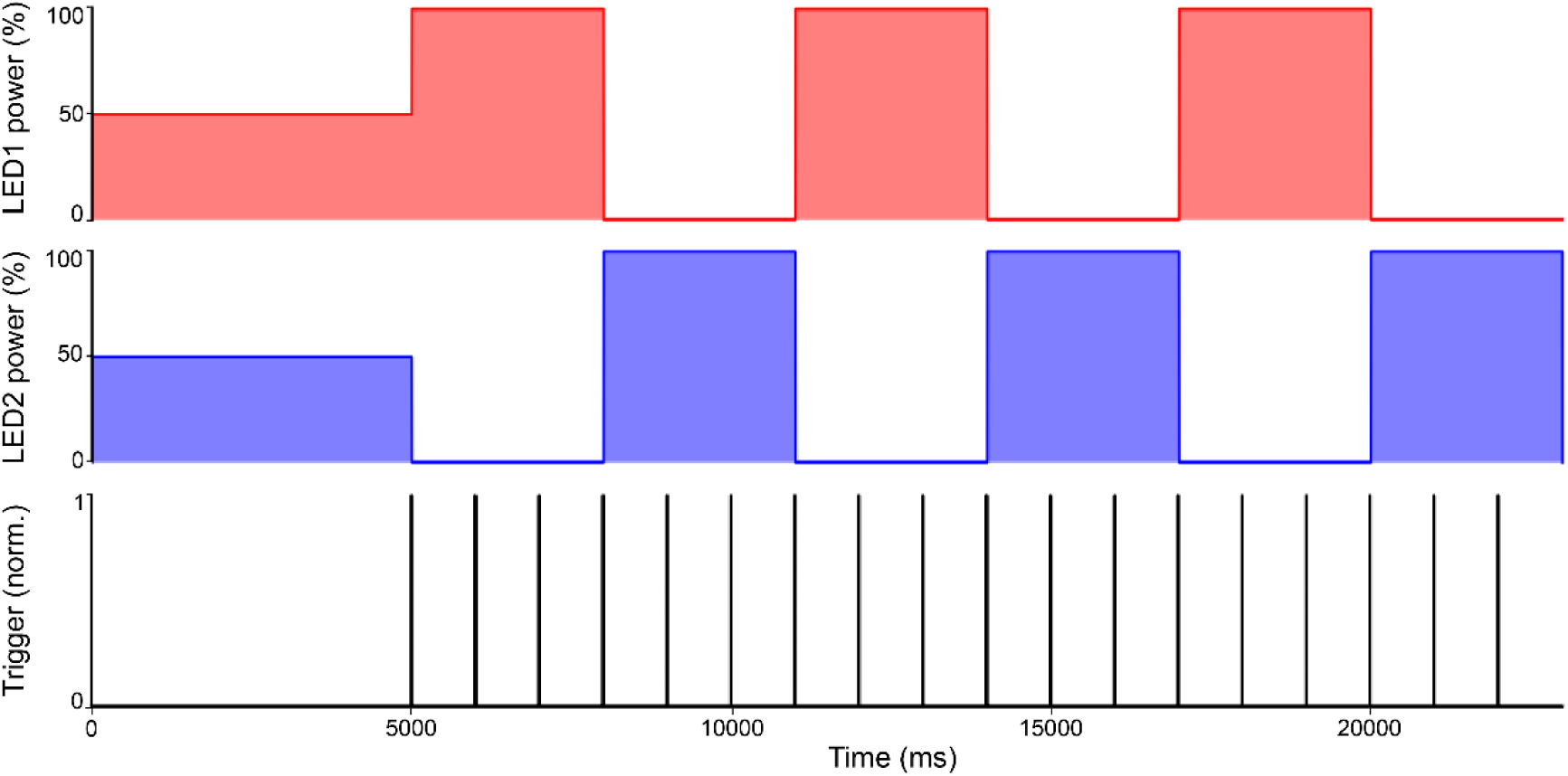
Stimulus example. LED sequence (On/Off steps of light over three loops) described above, along with trigger recording.

**Figure 8.**
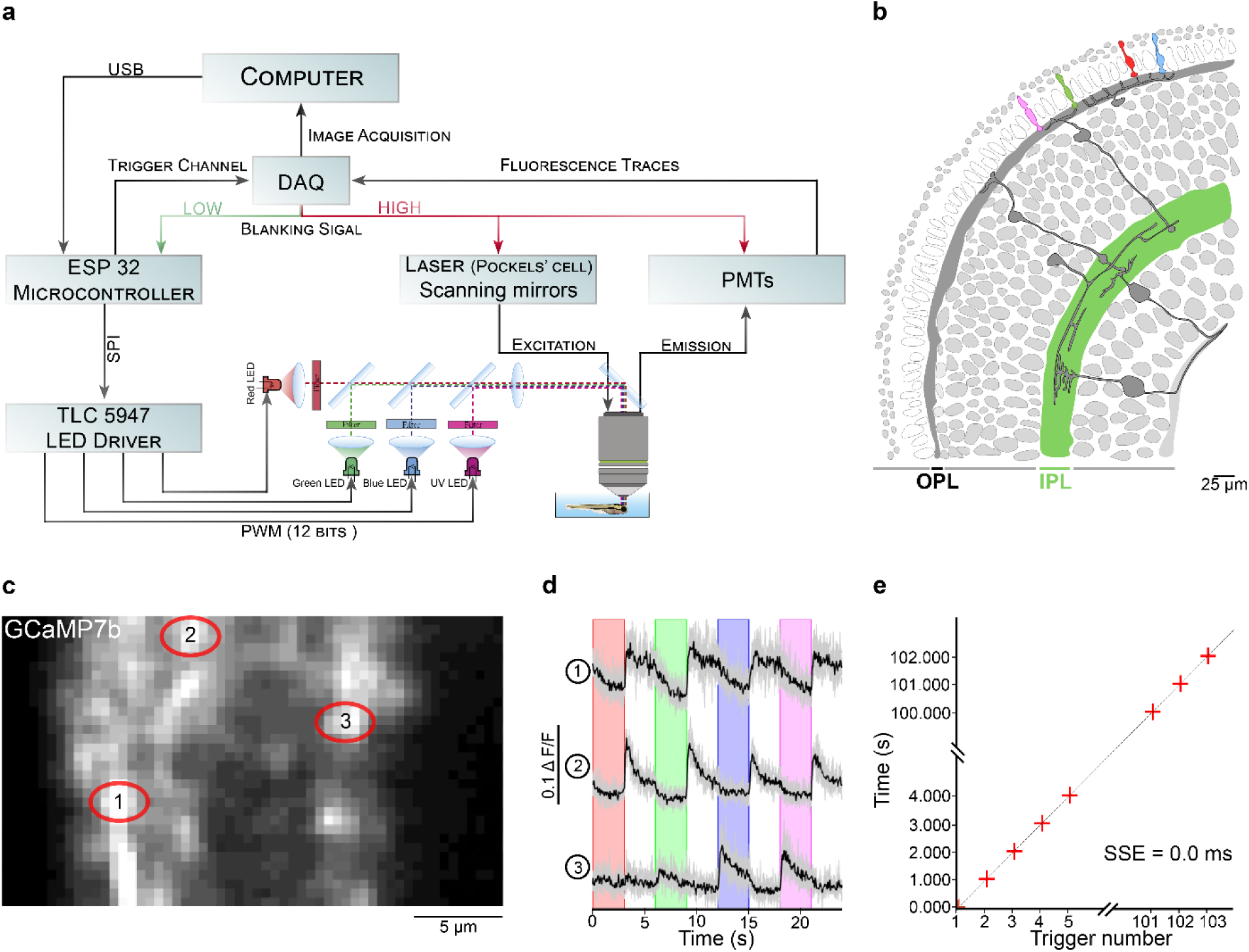
Zebrafish retina experiment. **a.** Overview of the setup described for the visual stimulation experiment performed on the tetrachromatic zebrafish. **b.** Drawing of the larval zebrafish retina highlighting the IPL. **c.** 2 photon scan field of the IPL with regions-of-interest marked by red circles. The 64×32 pixel image was obtained by at 1 ms scan rate. **d.** Ca^2+^ traces (mean traces in black, n=5 trials in grey) in response to consecutive red, green, blue and UV On/Off flashes. **e.** Trigger timing recorded by the DAQ highlighting its accuracy over time with a precision of 0.1 µs. **t(n+1) = t(n) + T**, where “t” is the recorded trigger time and “T” the trigger period.

### 2.5 *LED Zappelin’* without scan-synchronisation (optional)

Finally, our design can also be configured to work independent of an external blanking input, for example to function as a simple and time-precise LED controller. In this case, the ESP32 can be exchanged for a more economical Arduino Nano microcontroller. Since the Arduino Nano’s internal logic runs on 5V, while the ESP32 runs on 3.3V, we incorporated a second jumper at the bottom of the PCB to adjust the voltage depending on the microcontroller choice. A dedicated Arduino script is also provided to run the stimulator in this configuration.

## 3. Design files

### Design Files Summary

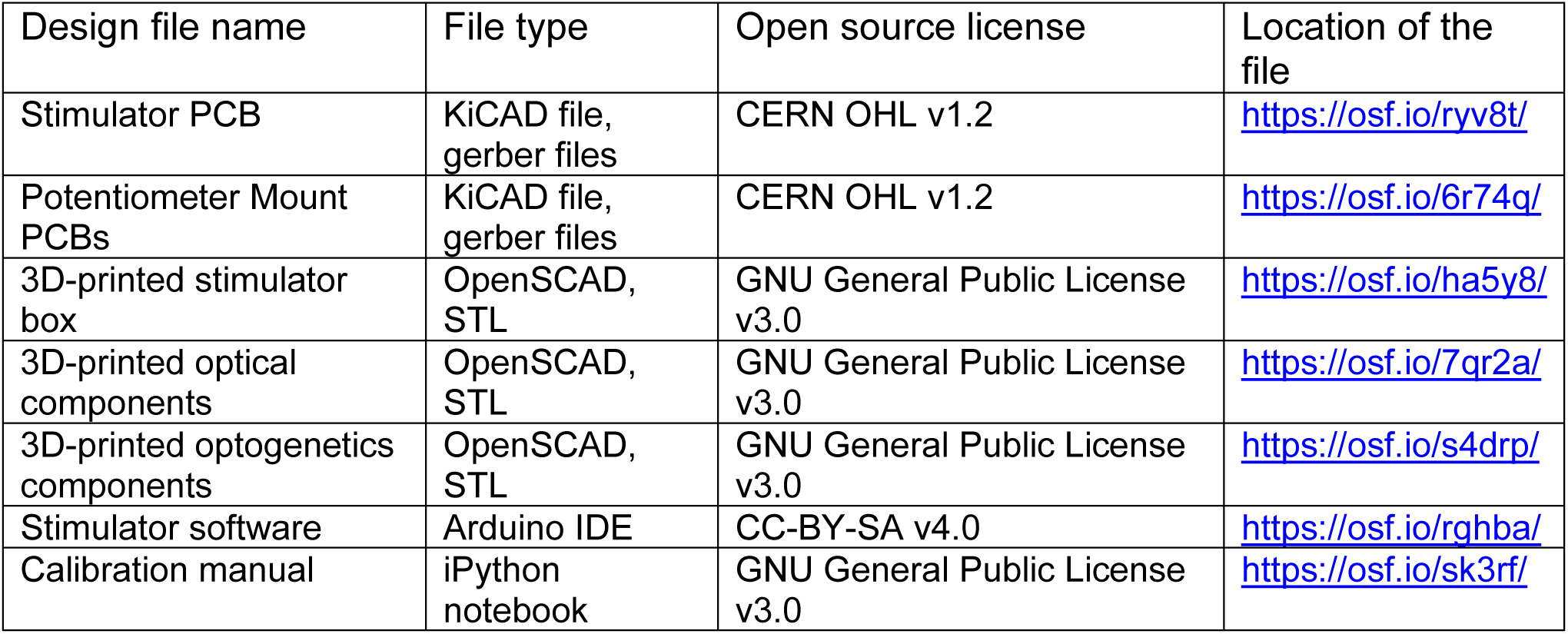

Readers interested in reproducing this system can use ready-to-order electronic design platform for open hardware projects: https://kitspace.org/, a PCB repository where all boards and components for them can be put in a “shopping cart” with one click.

#### Stimulator PCB

(Fig. 1b): This custom PCB is the core of the stimulator, to which each component is soldered.

#### Potentiometer mount PCBs

(Fig. 4): Optional PCBs that allow adding multi-turn trimmer potentiometers for fine tuning the LED supply current. We provide different versions for different numbers of LED channels (4, 8, 16 & 24).

#### 3D-printed stimulator box

(Fig. 5): Parts that fit and protect the electronics. The SCAD file is easily adaptable by the user.

#### 3D-printed optical components

(Fig. 6a): Parts used in combination with common optomechanical systems to hold LED, filter and dichroic mirrors in order to combine and collimate multiple spectral LEDs into a common light beam.

#### 3D-printed optogenetics components

(Fig. 6c-d): Parts that hold the sample and the stimulating LEDs. Chamber mounts are designed to fit a RC-40HP chamber (Thorlabs) and a 35mm Petri dish lid. A 3D-printed mounting platform fits the mounts onto a standard M6 rigid platform (Fig 9b).

**Figure 9.**
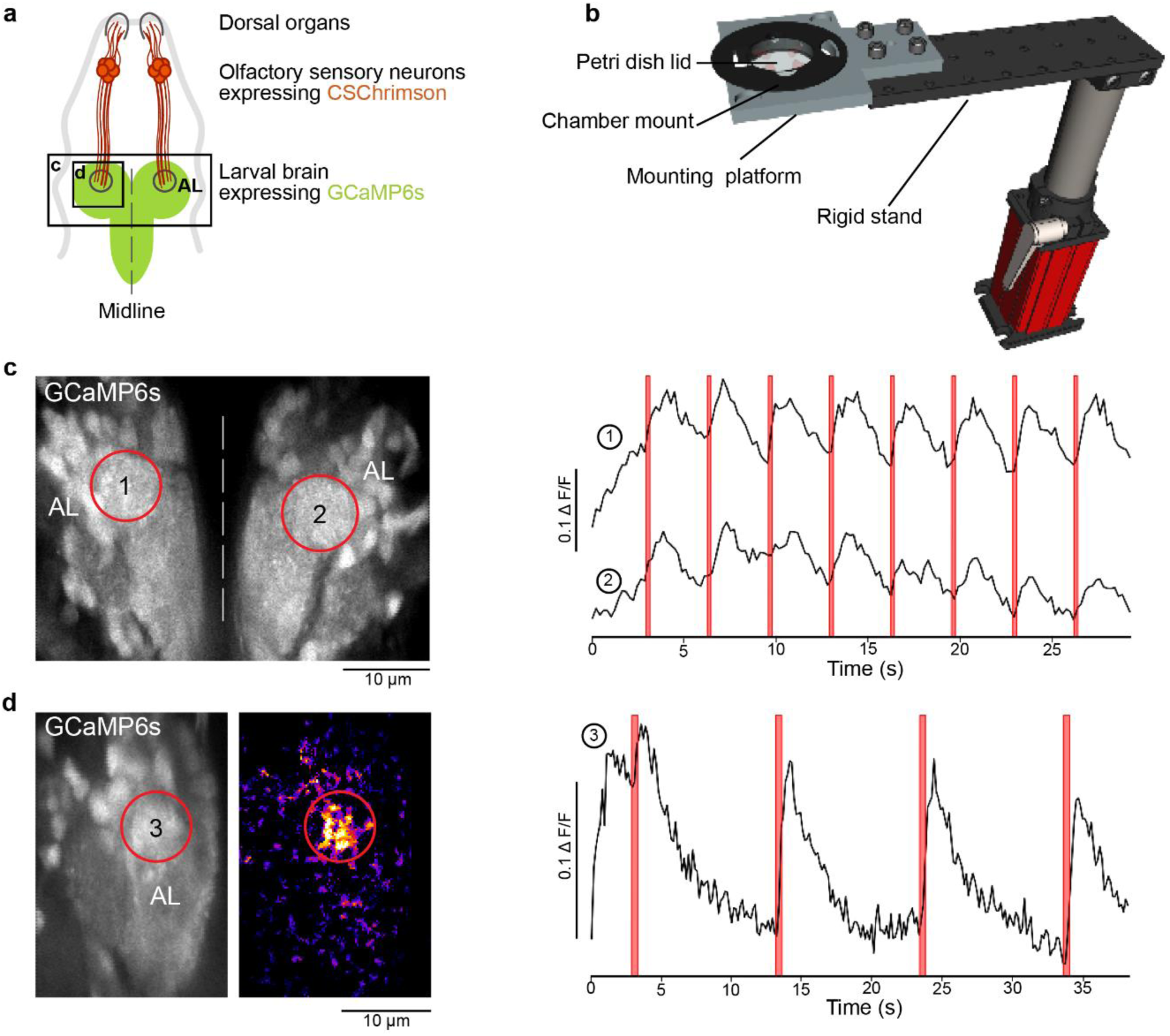
Drosophila optogenetics experiment. **a.** Schematic of a fruit fly first instar larval head expressing the red-shifted channel rhodopsin CsChrimson in olfactory sensory neurons and GCaMP6s in pan-neuronally. **b.** Rendering of the experimental setup: The mounting chamber (Fig. 6d) is placed in a 3D-printed holder (c.f. 3), screwed onto a rigid stand (ThorLabs). **c-d.** Standard deviation projections of 2 photon scan fields of the larval brain with antennal lobes marked by red circles (left) and Ca^2+^ traces in response to red flashes (right). **c.** Stimulation duration = 0.5 s, inter-stimulus interval = 3 s, image dimensions = 256 x 230, scan rate (lines) = 1,081 Hz, frame rate = 4.7 Hz. d. Stimulation duration = 0.5 s, inter-stimulus interval = 10 s, image dimensions = 256 x 170, scan rate (lines) = 1,077 Hz, frame rate = 6.34 Hz. Middle panel is a heatmap of pixel intensities showing high GCaMP6 fluorescence in the antennal lobe following optogenetic stimulation; obtained by subtracting a pre-stimulus from a during-stimulus image (median filter, kernel size = 2).

#### Stimulator Software

Arduino codes that control the stimulator. Easily modifiable to generate custom stimuli.

#### Calibration Manual

A Jupyter notebook (Python 3) with step by step instructions for intensity-calibrating the stimulator (an additional power-meter will be required).

## 4. Bill of Materials

| Description | Component | Number | Unit price | Total | Source of materials |
| --- | --- | --- | --- | --- | --- |
| PCBs | Stimulator + Potentiometer PCBs | 1 of each | ~\$10 | ~\$20 | JLCPCB |
| Microcontroller | Adafruit Huzzah32 – ESP32 Feather | 1 | \$19.95 | \$19.95 | Adafruit |
| LED Driver | Adafruit TLC5947 | 1 | \$14.95 | \$14.95 | Adafruit |
| BNC coaxial Connectors | Right angle BNC jacks | 2 | \$2.5 | \$5 | RS-components |
| 2-way JST connector | 2-way JST female connector housing | nLED | \$0.5 | nLED x \$0.5 | RS-components |
| 2-way JST PCB header | 2-way JST male connector PCB header | nLED | \$0.25 | nLED x \$0.25 | RS-components |
| 3-way JST connector | 3-way JST female connector housing | 1 | \$0.5 | \$0.5 | RS-components |
| 3-way JST PCB header | 3-way JST male connector PCB header | 1 | \$0.5 | \$0.5 | RS-components |
| Trimmer potentiometer | Multi-turn 10 kΩ through hole trimmer potentiometer | nLED | \$1.15 | nLED x \$1.15 | RS-components |
| Voltage Divider | 220 $\Omega$ through hole resistor | 1 | \$0.15 | \$0.15 | RS-components |
| Voltage Divider | 470 $\Omega$ through hole resistor | 1 | \$0.15 | \$0.15 | RS-components |
| Signal Inverter | 150 $\Omega$ through hole resistor | 1 | \$0.15 | \$0.15 | RS-components |
| Signal Inverter | 1 kΩ through hole resistor | 1 | \$0.13 | \$0.13 | RS-components |
| Power plug | 2.1 mm right angle DC socket | 1 | \$1.3 | \$1.3 | RS-components |
| PCB sockets | 2.54 mm pitch 16-way 1 row straight PCB socket | 6 | \$3.8 | \$22.8 | RS-components |
| Rocker LED switch | 3-pin LED Rocker ON/OFF SPST switch | 1 | \$5 | \$5 | RS-components |
| Total cost for 4 stimulating LEDs and 4 proxy LEDs stimulator | | | | \$105.78 | |

## 5. Build Instructions

### 5.1 Soldering the custom-designed PCB

The board is shown in Fig. 1b. For the microcontroller (left), two options are available for either the Arduino Nano (inner rows, no external line-synch option) or the ESP32 (outer rows, full stimulator). There is no need to solder more Japan Solderless Terminal (JST) pins beyond the number of LEDs required (here, we will show a version where 8 JST pins are connected: 4 for the stimulus LEDs, 4 for their proxy LEDs). On the right side of the board, the power plug and the 3-way JST header must be soldered into the respective sockets.

The jumper on the top (signal Inverter) allows the inversion of the Transistor-Transistor Logic (TTL) Blanking signals. For ScanImage users, the jumper should be placed in the upper position to switch off LEDs when the blanking signal is LOW. ScanM users should instead place it in the lower position to switch off LEDs when the blanking signal is HIGH.

The jumper at the bottom of the board (voltage divider) allows to set the voltage to 3.3 V (ESP32) or to 5 V (Arduino Nano). IMPORTANT: do not send 5 V signals to the ESP32. Since most TTL devices deliver 5 V pulses, we selected 220 Ω and 470 Ω resistors to bring an expected 5 V blanking signal to a 3.3 V input. Depending on the voltage range of the blanking signal, these resistors may need to be adjusted according to Ohm’s law: V1 + V2 = I*(R1+R2).

The Adafruit TLC5947 LED driver is configured by default to a current output level of 15 mA per channel, which is safe for nearly any standard LED. However, it is possible to operate at different currents by replacing the on-board resistor with a new through hole resistor. The driver can deliver up to 30 mA per channel (described in detail on the manufacturer’s datasheet: https://cdn-shop.adafruit.com/datasheets/tlc5947.pdf).

### 5.2 Mounting the potentiometers

To finely adjust each LED’s peak power, we added multiple-turn trimmer potentiometers to our design. To mount them, one simple solution is to manufacture the appropriate custom PCB (we provide multiple options for different numbers of channels). These extra PCBs fit tightly into the 3D-printed box. LEDs should be connected to the potentiometer in series as shown in Fig. 4.

### 5.3 Printing the Stimulator Box

We used OpenSCAD (freely available at www.openscad.org) to design the stimulator enclosure. The tolerance of the printer can be adjusted in the “USER Parameters” section of the script (tol = 0.1 mm by default, suitable for a reasonably well-calibrated Prusa i3 MK3 or Ultimaker 2). Each component can be displayed/designed individually in the “switches” section. Variables such as LED number (4 by default) and the potentiometer board dimensions can be adjusted in the “component parameters” section.

Default STL files can also be found on the repository and printed directly (4 stimulation LEDs + 4 proxy LEDs).

The PCB is mounted by adding 50mm M3 screws from the top via fitting holes in the stacking parts. The potentiometer board is fitted to the “back” part of the box, with trimmers fitting to their respective holes (Fig. 5b).

### 5.4 Mounting the proxy LEDs

The proxy LEDs provide convenient visualisation of the stimulus state for the experimenter. If this option is selected, 3 mm LEDs should be mounted at the back of the LED holder using 3 mm LED mounts and connected directly to their channel pins on the stimulator board (by default channels 5, 6, 7 & 8). As the LEDs are directly connected to an LED driver, no resistors are needed. Take note of their polarity (long LED leg should be connected to pin +). For aesthetics, cuttings of Teflon sheet or white paper (e.g.) can be placed in the LED holder slot to diffuse the LED light.

### 5.5 Mounting the stimulating LEDs

Each stimulating LED must be connected to its respective channel, taking note of their polarity (long LED leg should be connected to the positive pin).

For our visual stimulation setup, we combined all LED light sources into one beam which is projected through the objective to our model retina. We therefore constructed an optical cage system using a mixture of Thorlabs parts and 3D-printed objects to hold all filters and dichroic mirrors (Fig. 6a). This LED cage system was used and described in another publication (Franke et al., 2019).

For the optogenetics experiment we 3D-designed arenas where the sample sits, surrounded by four LEDs (Fig. 6c-d).

### 5.6 Connecting the stimulator to LEDs, the microscope’s DAQ and a computer

The stimulator can be externally powered anywhere between 5-30 V via the power port. Since the TLC5947 is a constant current LED driver, the voltage selection is not critical, however it should be slightly higher than the LED forward voltage (cf. LED driver datasheet). If desired, multiple LEDs can be connected to the same channel, however in this case the voltage supply must be adjusted accordingly (cf. LED driver datasheet).

For the standard line-synched stimulator version with an ESP32, a line-synched 5V TTL blanking signal BNC must be fed into the stimulator from the microscope’s DAQ (if the TTL is different from 5V, this can still be accepted provided the associated resistor is changed accordingly – see soldering paragraph). Note: Since for the default ESP32 version the blanking signal is used as the external clock, the stimulator will not execute any stimuli without it. If such line-synching is not required, consult the “simple” non-synchronised version that can be used with a simple Arduino Nano.

If required, connect the output trigger channel to the microscope’s DAQ. This signal generated by the stimulator by default sends a 3.3V pulse (if ESP32 is used, 5V for Arduino Nano) once at the start of the stimulus and then again, every 1,000 ms (1 Hz exactly). The trigger signal can for example be used to time-align acquired imaging data with the stimulus in postprocessing.

Finally, the board is connected to a computer via USB (micro USB for ESP32, mini USB for Arduino Nano).

## 6. Operation Instructions

### 6.1 Programming the ESP32 (or Arduino) on Windows systems

1. Download and install Arduino environment on the computer (www.arduino.org).
2. To use the ESP32, in addition:
  a. Install the latest SiLabs CP2104 driver (https://www.silabs.com/products/development-tools/software/usb-to-uart-bridge-vcp-drivers).
  b. Follow the installation instructions from the Espressif repository (https://github.com/espressif/arduino-esp32/blob/master/docs/arduino-ide/boards_manager.md).
3. Install the TLC5947 library
  a. Start Arduino and from the “Sketch” tab, select “Include Library” and open “Manage Libraries”
  b. From the search bar enter “TLC5947”
  c. Select and install the library
4. Open the Arduino script (2 versions available on the repository: *“2Photon_LED_Stimulator”* and “*Simple_LED_Stimulator”*, the second one being a simplified version of the first, independent of an external *blanking signal* input).
5. From the “Tools” tab:
  a. For the ESP32:
    i. Select from “Boards” the “Adafruit ESP32 Feather”.
    ii. From “Upload Speed”, select 921,600 (baud rate).
    iii. From “Flash Frequency”, select 80 Hz.
    iv. From “Port”, select the computer port to which the ESP32 is connected (if in doubt, unplug the board to see which ports are available, re-plug and observe which port is added). If the ESP is not recognised, check the driver installation (2a.), then check the micro USB cable (some USB cables do not work as not all their internal lines are connected).
  b. For the Arduino Nano
    i. Select from “Boards” the “Arduino Nano”
    ii. From “Processor”, select “ATmega328P” (option “Old Bootloader” for Arduino clones or older Arduino versions – if in doubt, try both)
    iii. From “Port”, select the computer port to which the Arduino is connected. If an Arduino clone is used, check that the proper driver is installed on the computer (consult its datasheet) and check the mini USB cable.
6. Compile and upload the code (clicking on the sideways arrow button on the top left).
7. The stimulator is ready to be used.

### 6.2 Operating the stimulator

The code is organised in five parts:

#### Stimulus Parameters

The code is designed to iteratively loop a pre-programmed stimulus sequence after an initial one-off optional preadaptation period.

– The number of loops is determined by “nLoops”. The stimulus will stop after finishing the n^th^ loop.
– IMPORTANT, the number of entries within the arrays must be the same and manually entered in “nArrayEntries” (including the pre-adaptation at position 1, see below).
– The “Scan_Logic” parameter corresponds to the x-mirror scan period in ms (i.e. = 2.0 if line speed is 2 ms per line and scan rate is 500Hz). This value must be changed if a different scan logic is used. This value defines the tempo of the entire stimulus (each time a blanking signal is counted, the code advances by an internal time-counter of Scan_Logic in milliseconds).
– The “array_LED#” arrays correspond to the stimulus sequence for each LED number. Here the number of arrays must be adjusted to the number of stimulating LEDs. All array positions will be read in synchrony, based on the timing array detailed below. IMPORTANT: The first entry is not part of the to-be-looped stimulus sequence, but instead defines the LED’s brightness for the preadaptation period. Note also that the first trigger output will start with the <u>second</u> entry (= the first entry of the stimulation sequence). The value entered at each position is the light intensity where 0 corresponds to no light and 100 to maximal light intensity. The value entered must range between 0 and 100.
– The “array_Time” array corresponds to the duration of each entry in ms; the first entry being the pre-adaptation that will only be played at the start of the stimulus, the sequence will then loop starting at the second position.

For example, if…

nLoops = 3;
nArrayEntries = 3;
Scan_Logic = 1.0;
Array_LED1 = {50,100,0};
Array_LED2 = {50,0,100};
Array_Time = {5000,3000,3000};

…the resultant stimulus will start with 5 seconds of both LEDs being set to 50% intensity (preadaptation) and will thereafter switch back and forth every 3 seconds between 100% and 0% power for the two LEDs in antiphase, for 3 repetitions (Fig. 7). Alongside, it will output one trigger signal every 1 s (fixed to this interval by default) once the looped portion of the stimulus starts (e.g. for later aligning the stimulus to imaging data). Throughout, the LEDs will be line-synched to a 1 ms scan logic.

#### Microcontroller Board Selection

Select if an ESP32 or an Arduino Nano is used.

#### Internal Definitions

This is the main definition part of the code which can be modified to:

- Add more LEDs than the 4 main and 4 proxy defined by default. (Global variables, the LED pins correspond to the pin number on the TLC5947).
- Adjust the trigger duration (25 ms by default).
- Adjust the trigger interval (1,000 ms by default).

#### Internal Methods

This is the main core of the code and should not be structurally changed (apart from adding more LEDs, as required).

#### Main Loop

This is where the serial user controls are defined. By default, when the serial monitor is open (magnifying glass on the top right corner in the Arduino IDE) and the baud rate at the bottom right of the window has been changed to 115,200. In this configuration, a manual command followed by pressing “enter” will trigger a stimulus:

By default:

- When “a + ENTER” is entered in the serial monitor, the stimulator will play the sequence with intensity scaled relative to the predefine “max1_LED#” powers (see below)
- When “b + ENTER” is entered, the same stimulus sequence will be played, but this time at the intensities defined by “max2_LED#” powers (see below)
- If “0 + ENTER” is entered during a stimulus sequence, all LEDs will be turned off and all loop counters will be reset.
- Further commands can easily be programmed by the user from the “Main Loop” part of the Arduino code.

It is important to note that the stimulation will only be played if a blanking signal is sent to the board.

### 6.3 Calibrating the stimulator

Stimulating LEDs can be approximately brought into a desired intensity regime by adding a serial resistor to limit the current they receive (c.f. 5.2). They can also be further calibrated within the code:

The TLC5947 is a 12-bit PWM grayscale driver, meaning that it offers up to 4,096 grey levels to adjust each LED power.

In the Arduino code there is a second tab called “LED_values” which hard-codes the maximum power an LED can get. Those values range from 0 (no current) to 4,095 (max current, 15 mA by default with potentiometer tuned all the way down, c.f. 5.1). In the default script we defined two distinct max values (max1 & max2) that can be called individually. The purpose here is to have the opportunity to use the same stimulus sequence at two different regimes of light intensities. More can be added manually by the user.

For the calibration, we suggest setting the max_LED# value to 4,095 (full power) and use successively a spectrometer and a power meter to adjust the LED brightness by finely turning the trimmer potentiometer at the back of the stimulator. As the LED output is linear relative to the values entered here (Fig. 3), any max_LED# value will be proportional to the LED power set up for the 4,095 value. The LED value (0-100%) entered in the stimulus sequence is linearly mapped to 0-max_LED#.

For a clear calibration procedure, we provide an easy step by step Jupyter notebook manual that can be found in our GitHub repository.

## 7. Validation and Characterization

### 7.1 Visual stimulation experiment

For colour vision experiments we recorded light-driven calcium signals under 2-photon from retinal bipolar cells *in vivo* in the tetrachromatic larval zebrafish (Zimmermann et al., 2018). We used a transgenic line expressing a genetically encoded biosensors for calcium at the bipolar cell synaptic terminals level *(ctbp2:SyGCaMP6)* (Dreosti et al., 2009) within the inner plexiform layer (IPL) (Fig. 8b).

Following established protocols (Euler et al., 2009), we used a Sutter-MOM microscope where light stimulation is displayed through the objective directly onto the fish retina along with the laser excitation. Fluorescence is also collected through the objective (Fig. 8a) as well as from below the stage (not shown).

We presented full field steps of red, green, blue and UV light to the fish eye (respectively 567, 480, 420 and 365 nm), and recorded evoked calcium signals as a readout of synaptic activity (Fig. 8c). We observed spectrally different tunings from distinct bipolar cell terminals (Fig. 8d) without detectable stimulus artefact across the scan.

In comparison to previous experiments performed on the same setup with a stimulator relying only on a basic microcontroller without LED driver (Zimmermann et al., 2018), LED-Zappelin completely (rather than “mostly”) eliminated the light-artefact on the sides of the scan. This is mostly due to the ESP32 processing power which also allows faster scanning rates compared to the performance achieved with traditional ATMega328 microcontrollers. Additionally, the use of an adequate LED driver providing linear 12-bit LED output (as opposed to, typically, 8-bit) has dramatically enriched possibilities in experimental design.

### 7.2 Optogenetics experiment

In addition to colour vision experiments, our LED Zappelin’ is well suited for optogenetic manipulation during 2-photon imaging. Here we illustrate this application from optogenetic circuit mapping in *Drosophila* larvae. Specifically, we recorded brain-wide calcium signals under 2P in response to optogenetic stimulation of all olfactory sensory neurons (OSNs). To this end, we expressed the red-shifted channel rhodopsin CsChrimson (Klapoetke et al., 2014) in OSNs and the genetically encoded calcium indicator GCaMP6s pan-neuronally (elav-Gal4;UAS-GCaMP6s/LexAOp-CsChrimson;Orco-LexA). We used first instar larvae that were fed from hatching on yeast paste supplemented with 0.4 mM all-trans retinal. Dissected larval heads with intact olfactory sensory organs (dorsal organs) and an exposed central brain (Fig. 9a) were immobilised in 3% low-melting-point agarose in physiological saline (Prieto-Godino et al., 2012) (in mM: 135 NaCl, 5 KCl, 5 CaCl2-2H2O, 4 MgCl2-6H2O, 5 TES (2-[[1,3-dihydroxy-2-(hydroxymethyl)propan-2-yl]amino]ethanesulfonic acid), 36 Sucrose, pH 7.15) and placed under the microscope in a 3D-printed chamber mount, itself placed on a 3D-printed stand fixed on a rigid stand (Fig. 9b, see also Fig. 6d). Red light stimulation was delivered from four sides of the recording chamber and GCaMP6s fluorescence intensity was collected by two detectors, one through the objective lens and a sub-stage PMT (not shown).

We presented full field illumination steps of 615 nm light lasting 0.5 s and an inter-stimulus interval of either 3 s (Fig. 9c) or 10 s (Fig. 9d). We observed robust stimulus-evoked activity in the primary olfactory sensory centres of the larval brain, the antennal lobes (AL, red outlines in Fig. 9c & d). As in our colour vision experiments (Fig. 8), we detected no light artefact at the sides of the scan, indicating near perfect time-synchronisation between the LEDs and the scan-lines. This continued being the case also during resonant scans (not shown).

## 8. Declaration of interest

AMC has a consultancy company providing services for Open Science: Chagas Science Consultancy, registered in the UK 12299826

## 9. Human and animal rights

All procedures were performed in accordance with the UK Animals (Scientific Procedures) act 1986 and approved by the animal welfare committee of the University of Sussex

## Supporting information

Supplementary Videos 1 and 2 in 3gp format

## Acknowledgements

We thank Thomas Euler for implementing an earlier version of a line-synched LED stimulator as e.g. used in (Baden et al., 2013), which heavily inspired the present design.

## Authors contribution

MJYZ conceived and implemented the stimulator with input from AMC, PB and TB. MJYZ performed the vision experiment on larval zebrafish. SP and LLPG performed the optogenetics experiments on *Drosophila*. MJYZ wrote the manuscript with help from TB and inputs from all authors.

## Funding

This work was supported by the European Research Council (ERC-StG “NeuroVisEco” 677687 to TB), Marie Sklodowska-Curie European Training network “Switchboard” (Switchboard receives funding from the European Union’s Horizon 2020 research and innovation programme under the Marie Sklodowska-Curie grant agreement No. 674901), the UKRI (BBSRC, BB/R014817/1 and MRC, MC_PC_15071 to TB), the Leverhulme Trust (PLP-2017-005 to TB), the Lister Institute for Preventive Medicine (to TB), the Marie Curie Sklodowska Actions individual fellowship (“ColourFish” 748716 to TY) from the European Union’s Horizon 2020 research and innovation programme. Research in the L.L.P.-G. is supported by The Francis Crick Institute and by the European Research Council (ERC StG (“EvoNeuroCircuit” 802531 to LLPG).

## Notes

https://github.com/BadenLab/LED-Zappelin

## References

Akerboom J, Chen TW, Wardill TJ, Tian L, Marvin JS, Mutlu S, Calderón NC, Esposti F, Borghuis BG, Sun XR, Gordus A, Orger MB, Portugues R, Engert F, Macklin JJ, Filosa A, Aggarwal A, Kerr RA, Takagi R, Kracun S, Shigetomi E, Khakh BS, Baier H, Lagnado L, Wang SSH, Bargmann CI, Kimmel BE, Jayaraman V, Svoboda K, Kim DS, Schreiter ER, Looger LL. 2012. Optimization of a GCaMP calcium indicator for neural activity imaging. J Neurosci. doi: 10.1523/JNEUROSCI.2601-12.2012

Alfonso-Garcia A, Li C, Bec J, Yankelevich D, Marcu L, Sherlock B. 2019. Fiber-based platform for synchronous imaging of endogenous and exogenous fluorescence of biological tissue. Opt Lett. doi: 10.1364/ol.44.003350

Baden T, Schubert T, Chang L Le, Wei T, Zaichuk M, Wissinger B, Euler T, Zaichuck M, Wissinger B, Euler T, Zaichuk M, Wissinger B, Euler T. 2013. A tale of two retinal domains: near-optimal sampling of achromatic contrasts in natural scenes through asymmetric photoreceptor distribution. Neuron 80:1206–17. doi: 10.1016/j.neuron.2013.09.030

Dreosti E, Odermatt B, Dorostkar MM, Lagnado L. 2009. A genetically encoded reporter of synaptic activity in vivo. Nat Methods 6:883–889. doi: 10.1038/nmeth.1399

Euler T, Franke K, Baden T. 2019a. Studying a light sensor with light: Multiphoton imaging in the retinaNeuromethods. doi: 10.1007/978-1-4939-9702-2_10

Euler T, Franke K, Baden T. 2019b. Studying a Light Sensor with Light: Multiphoton Imaging in the Retina. Preprints. doi: 10.20944/PREPRINTS201903.0244.V1

Euler T, Hausselt SE, Margolis DJ, Breuninger T, Castell X, Detwiler PB, Denk W. 2009. Eyecup scope-optical recordings of light stimulus-evoked fluorescence signals in the retina. Pflugers Arch Eur J Physiol 457:1393–1414. doi: 10.1007/s00424-008-0603-5

Franke K, Maia Chagas A, Zhao Z, Zimmermann MJ, Bartel P, Qiu Y, Szatko KP, Baden T, Euler T. 2019. An arbitrary-spectrum spatial visual stimulator for vision research. Elife 8. doi: 10.7554/eLife.48779

Helmchen F, Denk W. 2005. Deep tissue two-photon microscopy. Nat Methods. doi: 10.1038/nmeth818

Hohman B. 2007. The LED light source: A major advance in fluorescence microscopy. Am Biotechnol Lab. doi: 10.2345/0899-8205(2007)41[461:LLSMAI]2.0.CO;2

Icha J, Weber M, Waters JC, Norden C. 2017. Phototoxicity in live fluorescence microscopy, and how to avoid it. BioEssays. doi: 10.1002/bies.201700003

Klapoetke NC, Murata Y, Kim SS, Pulver SR, Birdsey-Benson A, Cho YK, Morimoto TK, Chuong AS, Carpenter EJ, Tian Z, Wang J, Xie Y, Yan Z, Zhang Y, Chow BY, Surek B, Melkonian M, Jayaraman V, Constantine-Paton M, Wong GKS, Boyden ES. 2014. Independent optical excitation of distinct neural populations. Nat Methods. doi: 10.1038/nmeth.2836

Marvin JS, Borghuis BG, Tian L, Cichon J, Harnett MT, Akerboom J, Gordus A, Renninger SL, Chen T-W, Bargmann CI, Orger MB, Schreiter ER, Demb JB, Gan W-B, Hires SA, Looger LL. 2013. An optimized fluorescent probe for visualizing glutamate neurotransmission. Nat Methods 10:162–170. doi: 10.1038/nmeth.2333

Mostany R, Miquelajauregui A, Shtrahman M, Portera-Cailliau C. 2014. Two-photon excitation microscopy and its applications in neuroscience. Methods Mol Biol 1251:25–42. doi: 10.1007/978-1-4939-2080-8_2

Pologruto TA, Sabatini BL, Svoboda K. 2003. ScanImage: Flexible software for operating laser scanning microscopes. Biomed Eng Online. doi: 10.1186/1475-925X-2-13

Prieto-Godino LL, Diegelmann S, Bate M. 2012. Embryonic Origin of Olfactory Circuitry in Drosophila: Contact and Activity-Mediated Interactions Pattern Connectivity in the Antennal Lobe. PLoS Biol. doi: 10.1371/journal.pbio.1001400

Svoboda K, Yasuda R. 2006. Principles of Two-Photon Excitation Microscopy and Its Applications to Neuroscience. Neuron. doi: 10.1016/j.neuron.2006.05.019

Yang W, Srivastava PK, Han S, Jing L, Tu CC, Chen SL. 2019. Optomechanical Time-Gated Fluorescence Imaging Using Long-Lived Silicon Quantum Dot Nanoparticles. Anal Chem. doi: 10.1021/acs.analchem.9b00517

Zimmermann MJY, Nevala NE, Yoshimatsu T, Osorio D, Nilsson DE, Berens P, Baden T. 2018. Zebrafish Differentially Process Color across Visual Space to Match Natural Scenes. Curr Biol. doi: 10.1016/j.cub.2018.04.075

